# Whole genome analysis of four Bangladeshi individuals

**DOI:** 10.1101/2020.05.21.109058

**Authors:** Salim Khan, Shahina Akter, Barna Goswami, Ahashan Habib, Tanjina Akhtar Banu, Carl Barton, Eshrar Osman, Samiruzzaman Samir, Farida Arjuman, Saam Hasan, Maqsud Hossain

**Affiliations:** Bangladesh Council of Scientific and Industrial Research, Dhaka, Bangladesh; Academica Solutions, London, United Kingdom; SciTech Consulting and Solutions, Dhaka, Bangladesh; National Institute of Cancer Research Hospital, Dhaka, Bangladesh; Dept. of Biochemistry and Microbiology, North South University, Dhaka, Bangladesh; NSU Genome Reseach Institute (NGRI), North South University

## Abstract

Whole-genome sequencing (WGS) is a comprehensive method for analysing entire genomes and this has been instrumental in characterizing the single nucleotide polymorphisms associated with different diseases including cancer, diabetes, cardiovascular diseases and many others. In this paper we undertake a pilot study for sequencing four Bangladeshi individuals and profiling their single nucleotide variants. Our findings shed possible light on specific biological pathways effected by such variants in this population.

## Main Text

Advancements in high-throughput next generation sequencing (NGS) technologies have enabled rapid and affordable sequencing of the whole human genome, allowing the generation of reference databases of population specific variants (1, 2).

The original Human Reference Genome had no representation from the subcontinent. Though subsequently, the 1000 genome project (3, 4) had some individuals of subcontinental origin, overall, the current database of human variants is still lacking in representation from this region. The Bangladeshi human genome needs to be analysed in greater depth and its population specific variants be added to the common variant databases so as to ensure they are a complete representation of the genetic variation observed in humans worldwide.

Bangladesh has a rapidly growing economy and in recent years has seen the establishment of a few state-of-the-art molecular biology research facilities such as the Genome Research Laboratory of Bangladesh Council of Scientific and Industrial Research (BCSIR). Here we undertake a pilot study using the whole genome sequences from four Bangladeshi individuals, labelled samples S1, S6, S19 and S21, to gain the first understanding of the single nucleotide variations (SNVs) that are unique to this population. The primary goal of this study was to identify unique variant-bearing genes in Bangladeshi individuals and to subsequently analyse their phenotypic impact in terms of gene functions and diseases.

The initial sequencing and mapping provided between 1.1 billion to 1.46 billion reads. Samples S1, S6 and S19 produced 1.3 to 1.46 billion reads, while the reads for S21 dropped down to around 1.1 billion. For each of the samples, under 25% of all reads were unmapped. The total alignments were around 1387640908, 1498505945, 1387640908, 1023927992 for samples S1, S6, S19 and S21, respectively (Table 1).

**Table 1:**
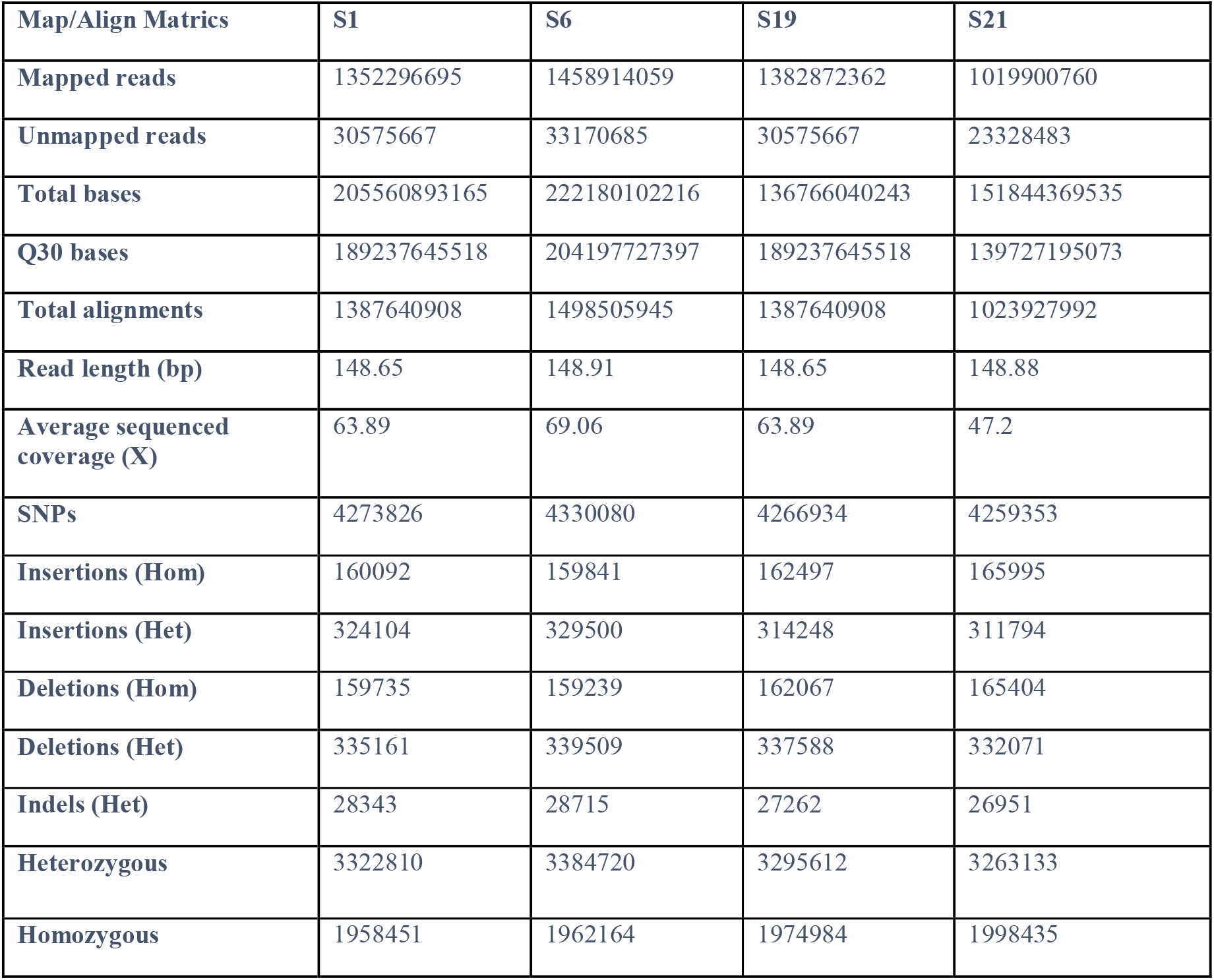
Summary of whole genome sequencing results for all four samples.

| Map/Align Matrics | S1 | S6 | S19 | S21 |
| --- | --- | --- | --- | --- |
| Mapped reads | 1352296695 | 1458914059 | 1382872362 | 1019900760 |
| Unmapped reads | 30575667 | 33170685 | 30575667 | 23328483 |
| Total bases | 205560893165 | 222180102216 | 136766040243 | 151844369535 |
| Q30 bases | 189237645518 | 204197727397 | 189237645518 | 139727195073 |
| Total alignments | 1387640908 | 1498505945 | 1387640908 | 1023927992 |
| Read length (bp) | 148.65 | 148.91 | 148.65 | 148.88 |
| Average sequenced coverage (X) | 63.89 | 69.06 | 63.89 | 47.2 |
| SNPs | 4273826 | 4330080 | 4266934 | 4259353 |
| Insertions (Hom) | 160092 | 159841 | 162497 | 165995 |
| Insertions (Het) | 324104 | 329500 | 314248 | 311794 |
| Deletions (Hom) | 159735 | 159239 | 162067 | 165404 |
| Deletions (Het) | 335161 | 339509 | 337588 | 332071 |
| Indels (Het) | 28343 | 28715 | 27262 | 26951 |
| Heterozygous | 3322810 | 3384720 | 3295612 | 3263133 |
| Homozygous | 1958451 | 1962164 | 1974984 | 1998435 |

After the variant calls, all four samples contained between 5 million and 5.5 million variants. Sample S1 gave us 5,279,748 variants and after removing variants with QUAL <20 total number of variants were found 5,000,704. Sample S6 had 5,345,421 variants initially and 5,064,885 after removing low quality calls. Sample S19 produced 5,269,076 variants with low quality calls and 4,970,655 calls without them. Lastly Sample S21 produced 5,260,335 calls with low quality variants and 4,966,352 variants after they had been filtered. After filtering out the common variants, approximately 900,000 variants were removed from each of the datasets. We observed that the number of variants per chromosome correlated with size. Chromosome 1 had the most number of variants, averaging around 4.13 million for four samples, while Chromosome 22 had the least, averaging around 80,500. The protein coding genes with the highest numbers of variants were *EMBP1, TTTY23, HLA, ACTR3BP2, ACTR3BP5, ALG10, XLOC, EPHA3, CWH43, CSMD1, HCN1, SLC25A51P1 and FRG1CP*. We found an identical number of variants within these genes for all four of the individuals. EMBP1 has 13, 820 SNP variants identified in the NCBI database, here we found 18,020. ACTR3BP2 has 1467 variants listed in the database, we found 11,458. TTTY23 had the most significant deviation from other databases. NCBI SNP lists the gene as containing 7 known variants, we found 13,030 (11). A list is shown in Table 2 in which we have shown the top 22 genes with most variants.

**Table 2:**
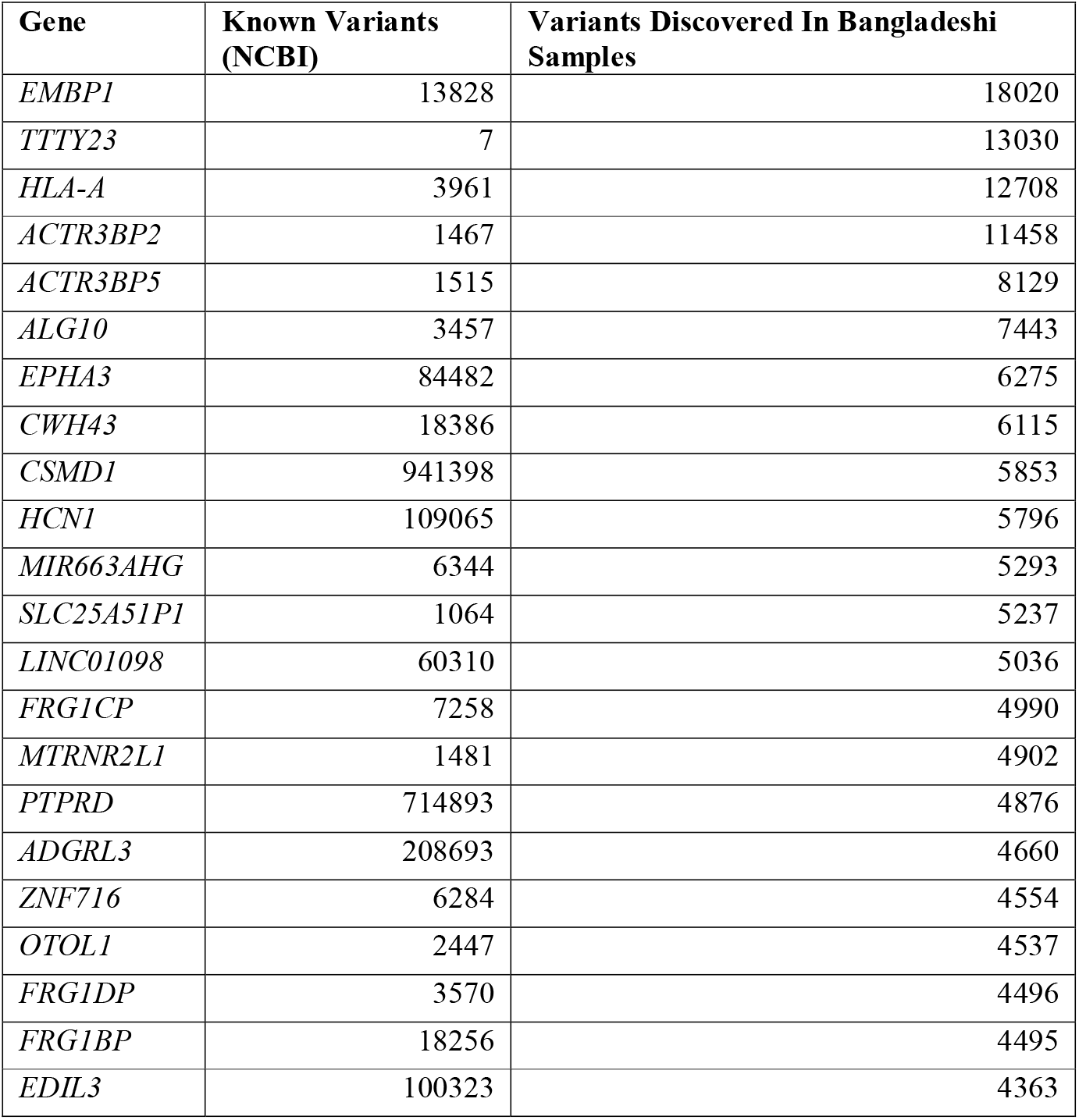
The protein coding genes with the highest number of variants among the Bangladeshi individuals.

In terms of exonic variants, all four samples produced 25,000 variants that occurred within exonic regions of genes. Among them, 11,582 were nonsynonymous mutations, 12,296 were synonymous, 192 were nonframeshift insertions, 218 were nonframeshift deletions, 110 were stop-gain variants, 116 were frameshift deletions, 98 were frameshift insertions, 10 were stop-loss and 378 were unknown. Out of all the exonic variants, 9524 were homozygous variants.

Examples of variants we found that led to protein coding changes include the *ABC12* variant at Chr7:48279214 that causes a change from Arginine to Tryptophan at position 2674 of the encoded protein, the *A2M* gene nonsynonymous variant at chr12:9095637 that causes a change from Asparagine to Aspartic Acid at position 639 of the protein, and others. Although none of these were unique to the Bangladeshi individuals.

On the other hand, a number of genes produced far fewer variants than what may have otherwise been expected. *RBFOX1* for example has 949,641 variants associated with it, we were only able to find 2375. *CNTNAP2* has 572,590 variants associated with it. Among them, we were only have to discover 319 in our samples.

After the ontology analysis using DAVID, we considered the functional annotation chart which listed the functions that could be affected by the variant containing genes. The P-Value and Benjamini scores indicate the significance of the genes in question to the respective pathways. Table 3 shows the diseases implicated in the ontology results for all four samples. In particular, we found heart related diseases and body fat and mass associated disorders to be implicated.

**Table 3:** Functions effected by highly variable genes in the Bangladeshi individuals.

| Associated Function | P-Value | P-Value | Benjamini Value |
| --- | --- | --- | --- |
| Cholesterol, HDL | 18.2 | 3.70E-03 | 7.80E-01 |
| Glucose | 13.6 | 1.40E-02 | 9.40E-01 |
| Respiratory Function Tests | 13.6 | 1.50E-02 | 8.70E-01 |
| macular degeneration | 13.6 | 1.50E-02 | 7.90E-01 |
| ADHD | 9.1 | 3.00E-02 | 9.20E-01 |
| Echocardiography | 13.6 | 3.40E-02 | 9.00E-01 |
| Mucocutaneous Lymph Node Syndrome | 9.1 | 3.80E-02 | 8.90E-01 |
| Triglycerides | 13.6 | 4.30E-02 | 8.90E-01 |
| Asthma | 13.6 | 4.70E-02 | 8.80E-01 |
| Body Mass Index | 13.6 | 5.00E-02 | 8.80E-01 |
| hepatitis C, chronic | 9.1 | 6.20E-02 | 9.10E-01 |
| Forced Expiratory Volume | 9.1 | 6.40E-02 | 8.90E-01 |
| Body Fat Distribution | 9.1 | 7.70E-02 | 9.20E-01 |
| Hematocrit | 9.1 | 8.40E-02 | 9.20E-01 |
| Behcet Syndrome | 9.1 | 9.70E-02 | 9.40E-01 |
| Lipoproteins | 9.1 | 9.90E-02 | 9.30E-01 |

With regards to disease associated SNPs, our samples contained the chr6:121447564 G>A variant that has been implicated in heart complications. Although the chances of its pathogenicity have been described as benign (ClinVar accession: VCV000137482) (14).

Here we report the whole genome sequencing of four Bangladeshi individuals carried out at Genome Research Laboratory of BCSIR, Bangladesh. One of the major objectives was to set up the baseline genomic catalogues of Bangladeshi population. As a result of this study we found around 900,000 variants previously identified in other genomes as well as nearly 5 million unique variants within the Bangladeshi genomes that could have possible functional implications. That number is expected come down following the continuation of these studies on larger sample sizes and more in-depth statistical validation. A number of genes containing significant numbers of variants from the Bangladeshi samples implicated various heart associated disorders when analysed. Genes *CSMD1, EDIL3, EPHA3, OTOL1, PTPRD* and *ZNF16* were all linked with heart or heart associated disorders by DAVID’s algorithm (5, 8). Previous studies have often listed cardiovascular disease as one of biggest causes of mortality in Bangladesh (7) and possible genetic links that may predispose the population to these conditions should be investigated further. Other effected functions include ADHD, glucose metabolism, and respiratory functions (table 3). All of these pathways returned enrichment scores that were well below the 0.05 threshold. The main limitation of this study was the smaller sample size. A more extensive sampling and sequencing of the population is being carried out at Genome Research Laboratory, BCSIR and the baseline study would provide data for establishing a more acceptable SNP map of the Bangladeshi human genome.

## Methods

### Sequencing

This study was approved by the Ethical Committee under the National Institute of Cancer Research and Hospital, Mohakhali, Dhaka-1212, (Ref.No.NICRH/Ethics/2019/525, Date: 22.09.2019) Bangladesh which is consistent with the declaration of Helsinki-Ethical Principles, October 2008. All the participating members provided informed written consent consistent with the experiment. A small aliquot (~5ml) of blood sample was collected from each individual and genomic DNA was extracted by using Maxwell RSC whole blood DNA extraction kit (Promega) according to the manufacturer’s instructions. The 300 ng gDNA of all four samples were used to prepare paired-end libraries with the Nextera ™ DNA Flex Library Preparation kit with an average insert size of 600 bp for all four samples according to the manufacturer’s instructions (Illumina Inc., San Diego, CA).

### Variant Calling, Annotation and Analysis

Illumina Basespace Sequence hub, Dragon Germline 3.4.5 (DRAGEN Host Software Version 05.021.332.3.4.5 and Bio-IT Processor Version 0×04261818) was used for mapping and variant calling. The VCF files were annotated using Annovar (12). As we wanted to identify variants unique to the Bangladeshi population, common variants known to occur in other populations were removed. For this purpose, we obtained the common human variant datasets from the UCSC and NCBI repositories (9, 11). These common variants were subsequently removed from our samples using bedtools (10) and some manual filtering using R. Finally, we shortlisted the top 1000 genes with the highest numbers of variants and carried out a Gene ontology was done using DAVID (8).

### Data Availability

The data supporting the conclusions of this article are included within the article. Raw Sequence data for four samples are available under the SRA accession number: PRJNA606337.

## Declaration

### Author Contributions

SK, SA, AH, TB and BG are participated in performing the experiment. SA, TA, BG, and SA carried out data analyses. CB, SS and EO assisted in developing pipelines and designing the workflow. FA carried out clinical examinations of subjects. SK, SH and MH wrote the manuscript. SK and MH conceived and oversaw the study. All authors read and approved the manuscript.

### Consent for publication

Authors have agreed to submit it in its current form for consideration for publication in the journal.

### Ethics approval and consent to participate

This research work has been carried out after complied with the national laws and regulations of the country and “WMA declaration of Helsinki-Ethical Principles for Medical Research Involving Human Subjects, amended ethically approved by National Institute of Cancer Research and Hospital. No. NICRH/Ethics/2019/525.

